# Cytokine and Immunoglobulin Dynamics in Phage Therapy: Insights from Clinical Cases

**DOI:** 10.1101/2025.11.21.689765

**Authors:** Samantha R. Ritter, Justin Clark, Keiko Salazar, Saima Aslam, Anthony Maresso

**Author notes:** Correspondence (S.A.); (A.M.). Joint senior authors (Saima Aslam, Anthony Maresso).

## Abstract

**Background:** Bacteriophage therapy is increasingly used for antibiotic-refractory infections, yet its immunologic effects in humans remain poorly defined. We evaluated cytokine and immunoglobulin responses in nine compassionate-use cases of intravenous phage therapy, including solid organ transplant recipients and patients with device-associated or refractory bacterial infections.

**Methods:** Serial serum samples were analyzed by multiplex Luminex immunology and isotyping assays, with results correlated to host immune status, infecting organism, and clinical outcome.

**Results:** Cytokine patterns were heterogeneous and appeared to reflect the infecting pathogen more than phage exposure. *Pseudomonas aeruginosa* infections elicited broad pro-inflammatory cytokine increases (notably MCP-3 and TNF-β), while *Staphylococcus aureus* infection was associated with overall cytokine reduction except for IL-6. Most immunocompetent patients exhibited an early IgM response within 1–2 weeks of phage initiation, whereas immunocompromised hosts demonstrated attenuated antibody levels. Neither cytokine nor humoral responses correlated with clinical outcome. In one case, serum neutralization developed against a specific phage but not against a subsequent, distinct phage cocktail targeting the same organism, suggesting variability in phage immunogenicity.

**Conclusion:** Overall, intravenous phage therapy frequently induces IgM responses, which are more pronounced in immunocompetent patients, while cytokine dynamics depend more on pathogen and immune status than on phage administration itself. These findings underscore the need for standardized immune profiling in clinical phage trials to delineate beneficial versus detrimental immune responses and to inform the design of phage formulations with optimized immunogenic profiles.

## Introduction

Phage therapy has been used to treat antibiotic-refractory infections with variable success, and clinical trials are ongoing.^1,2^ The immune response to phages—particularly serum neutralization seen in both animal models and humans—remains poorly defined. Murine studies with *Pseudomonas* phages suggest that bacterial clearance may result from combined phage and host immune activity, but most work has been limited to animal models, leaving major gaps in human data.^3^ We sought to characterize immune responses before, during, and after phage therapy in recent compassionate-use cases, focusing on cytokine and immunoglobulin patterns associated with clinical outcomes and the potential influence of immunosuppression.

## Materials and Methods

Serum samples were collected from patients undergoing compassionate use phage therapy at the University of California San Diego (UCSD) under local Institutional Review Board (IRB) protocol #200163 after obtaining informed consent. Each patient received phage therapy under individual, expanded access Investigational New Drug (IND) applications with oversight from the US Food and Drug Administration (FDA) and local IRB.

Patient serum samples were analyzed using multiplex Luminex immunology and isotyping assays, with results normalized to baseline and log-transformed for visualization as heatmaps. Raw assay data were integrated with de-identified clinical metadata (infection type, pathogen, treatment timeline, and outcome) and assessed using AI-assisted pattern recognition; trends were subsequently validated in GraphPad Prism (v10.4.1). Serum neutralization assays were performed by incubating patient sera with the phage cocktail and enumerating plaques to determine titers. Cytokine and immunoglobulin responses were grouped by treatment timepoints (baseline, Weeks 1–4, and post-treatment), and statistical comparisons were performed using Mann-Whitney U tests with Holm-Šídák correction for multiple comparisons. Full assay protocols, preprocessing steps, and instrument specifications are provided in the Supplement.

## Results

Seven patients received nine separate courses of compassionate-use phage therapy under FDA/IRB oversight, including three solid organ transplant recipients and three with device infections.^1,4–7^ Three were treated with phage manufactured at TAILOR Labs. Most infections were due to Gram-negative rods (3 *Escherichia coli*, 4 *Pseudomonas aeruginosa*, 1 *Klebsiella pneumoniae*), with one *Staphylococcus aureus* case (Table S1).

Across the cohort, cytokine responses varied considerably (Figure 1–2). Immunocompetent hosts demonstrated consistently higher levels of IL-6, TNF-α, IL-10, and IL-1β compared with immunosuppressed patients, though these differences were not statistically significant. No consistent association was observed between cytokine levels and clinical outcome (Figure 3).

**Figure 1.**
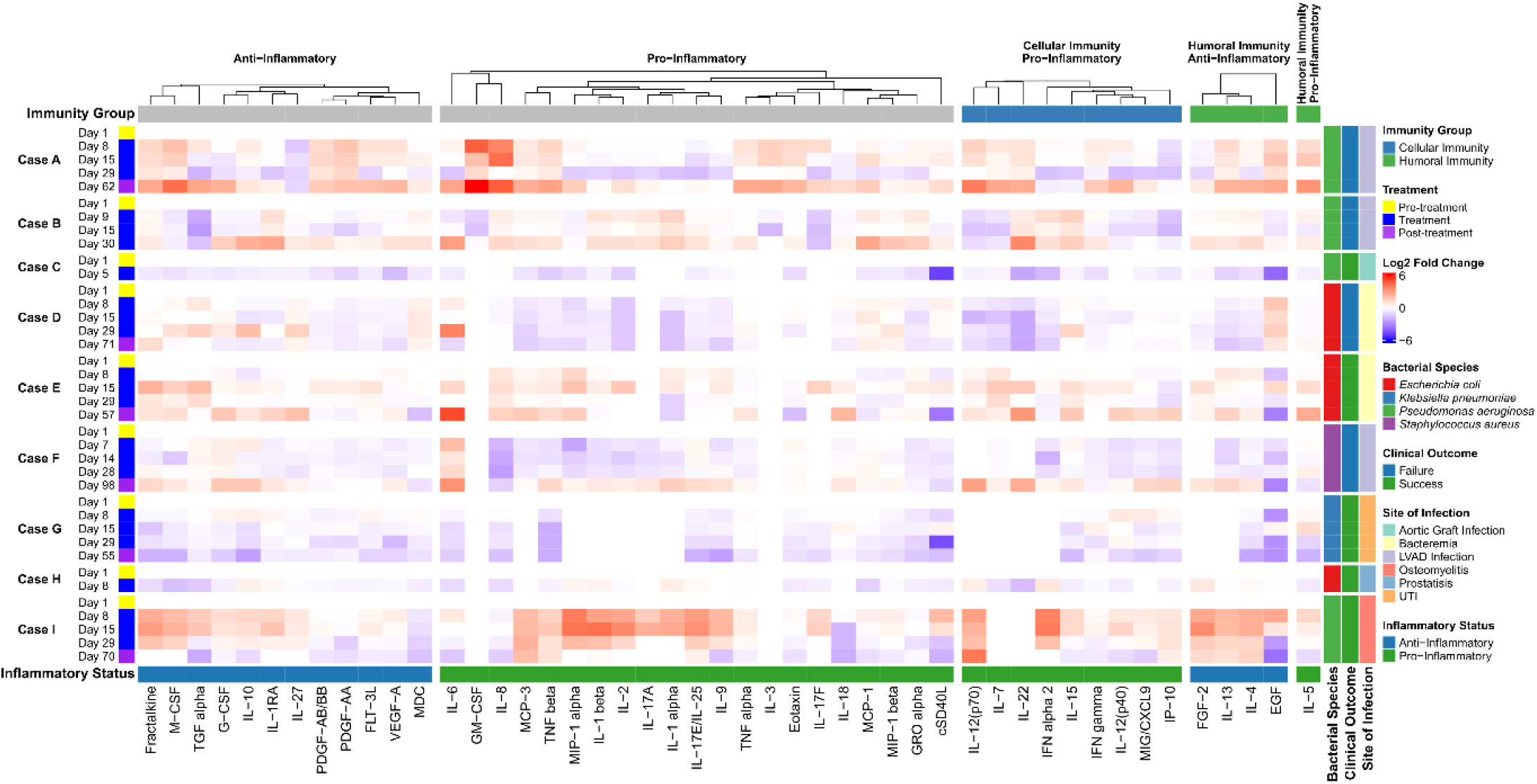
Unified heatmap of patient cytokine levels throughout the course of treatment. This heatmap groups cytokines into four categories; anti-inflammatory, pro-inflammatory, cellular immunity pro-inflammatory, and humoral immunity anti-inflammatory. Additional metadata is displayed to the right of the heatmap with a legend.

**Figure 2.**
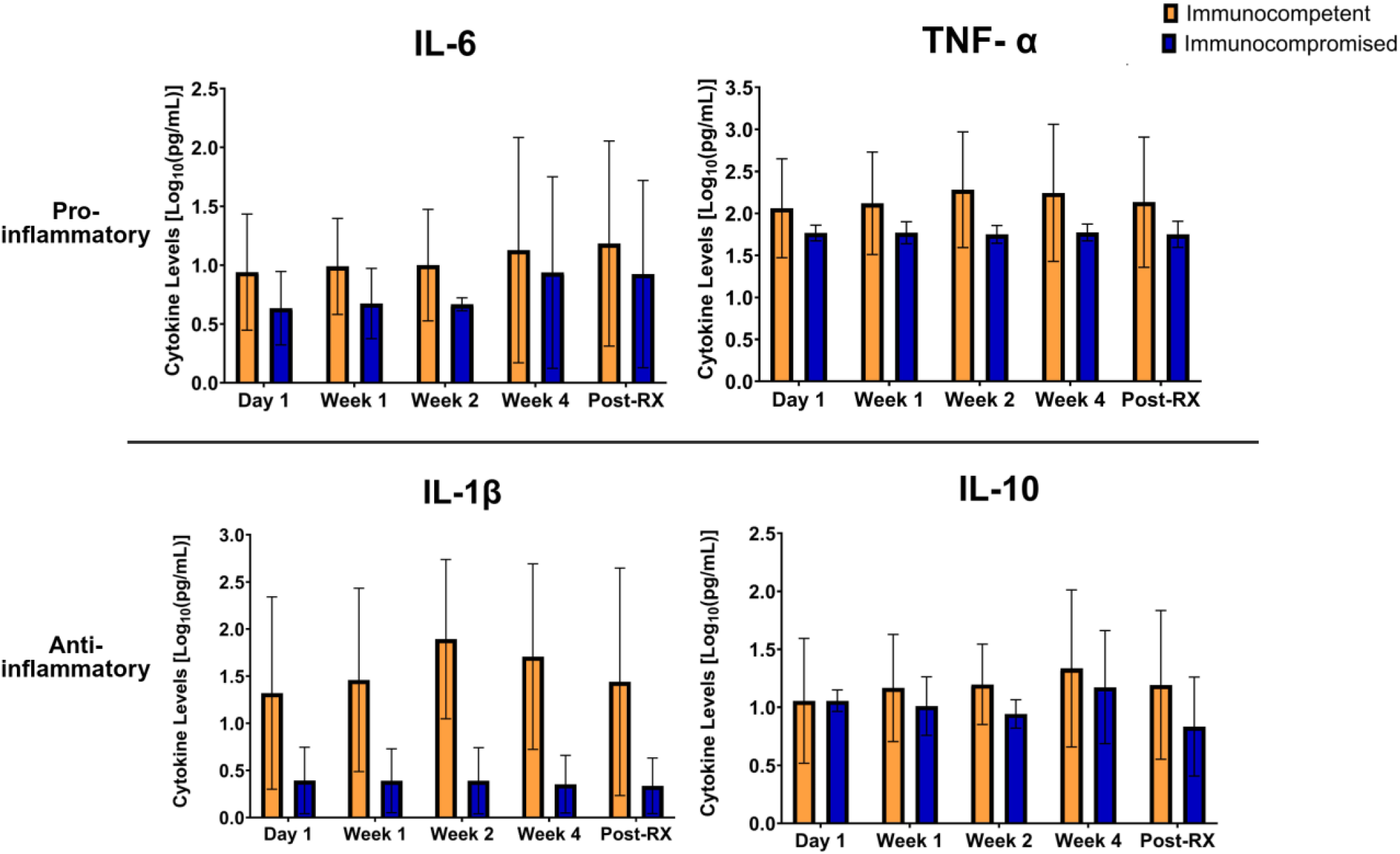
Grouped bar graph showing the difference of cytokines levels (pg/mL) transformed to Log_10,_ at different timepoints of treatment. Cytokine levels were measured from patient serum samples and were grouped into immunocompetent and immunocompromised groups. Cytokine levels were averaged for each group at every timepoint. The x-axis contains the timepoints in treatment in which cytokine levels were measured. Differences between the immunocompetent and immunocompromised groups were not found to be significant for all cytokines.

**Figure 3.**
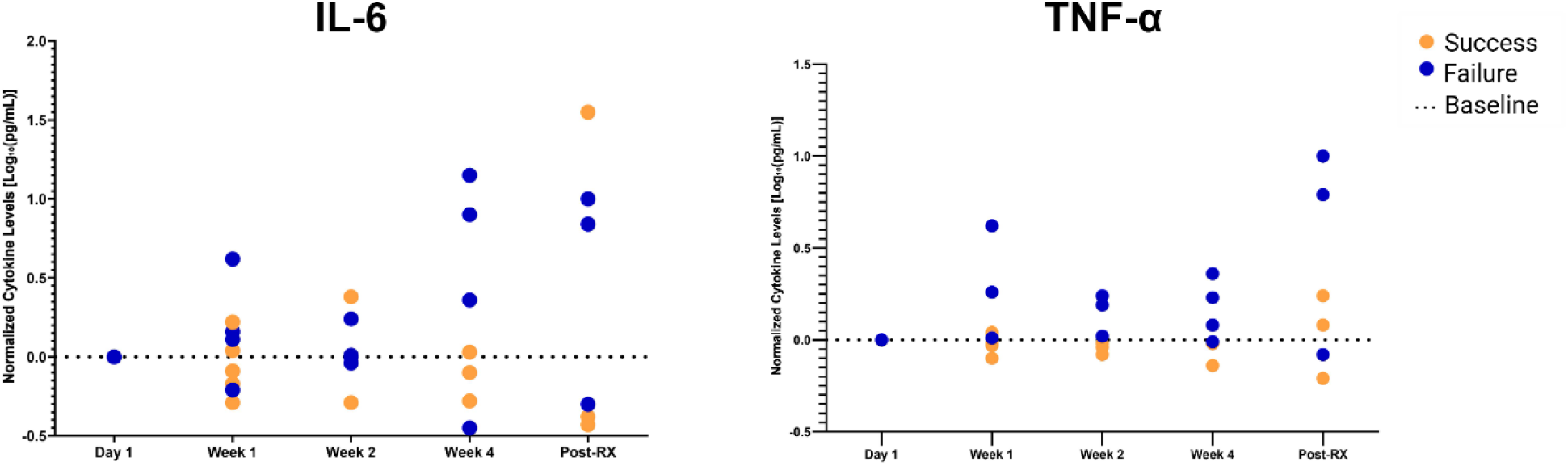
Plot of IL-6 and TNF-α levels throughout the course of treatment. For each separate case, data was normalized to the baseline value (day 1) by subtracting the baseline value from all other time points. Data was then transformed using Log_10_. Orange indicates success, and blue indicates failure. The x-axis contains the timepoints in treatment in which cytokine levels were measured. Differences between the success and failure groups were not found to be significant for both cytokines.

Cytokine responses appeared organism specific. In the *S. aureus* case, most pro-inflammatory cytokines decreased during phage therapy except IL-6, which rose; in contrast, *P. aeruginosa* cases demonstrated broad increases in pro-inflammatory cytokines, notably MCP-3 and TNF-β (Figure 4). This may indicate that the infecting organism is a stronger driver of cytokine response than the use of phage therapy.

**Figure 4.**
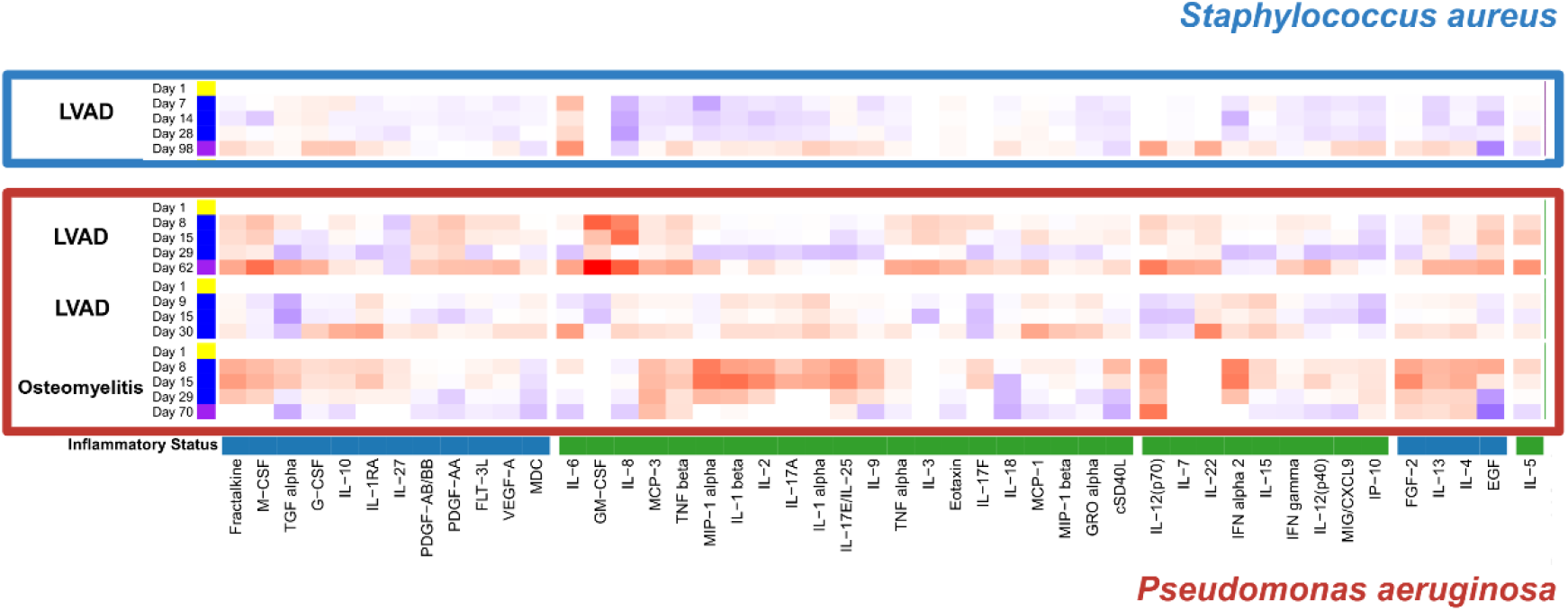
Cytokine response in patients with left ventricular device infection due to *Staphylococcus aureus* (n=1, blue) compared to *Pseudomonas aeruginosa* (n=3, red). The top two patients in the red box had an infected LVAD, and the patient on the bottom had pelvic osteomyelitis. Red indicates an increase in cytokine levels and blue indicates a decrease in cytokine levels.

As noted in Figure 5, seven of nine phage therapy cases developed an IgM response within 1-2 weeks after initiation of intravenous (IV) phage; the majority of these were in immunocompetent individuals. Levels of each immunoglobulin isotype varied per individual. Isotype levels in immunocompetent hosts were higher when compared to those that were immunocompromised (Figure 6). IgM had a higher median at all measured time points for immunocompetent hosts; Day 1–1·7 times higher (IQR 172·4-14·5), Week 1–2·4 times higher (IQR 181·9-30·3), Week 2– 5·0 times higher (IQR 213·8-126·6), Week 4–4·3 times higher (IQR 171·0-104·1), Post-RX–28 times higher (IQR 129·3-62·2). There was a peak in IgM response during the first or second week of treatment with phage therapy. In contrast, in the immunocompromised patients, there was little to no increase in IgM throughout the entire course of treatment. Additionally, all immunoglobulin isotypes in the immunocompromised hosts decreased over the course of treatment.

**Figure 5.**
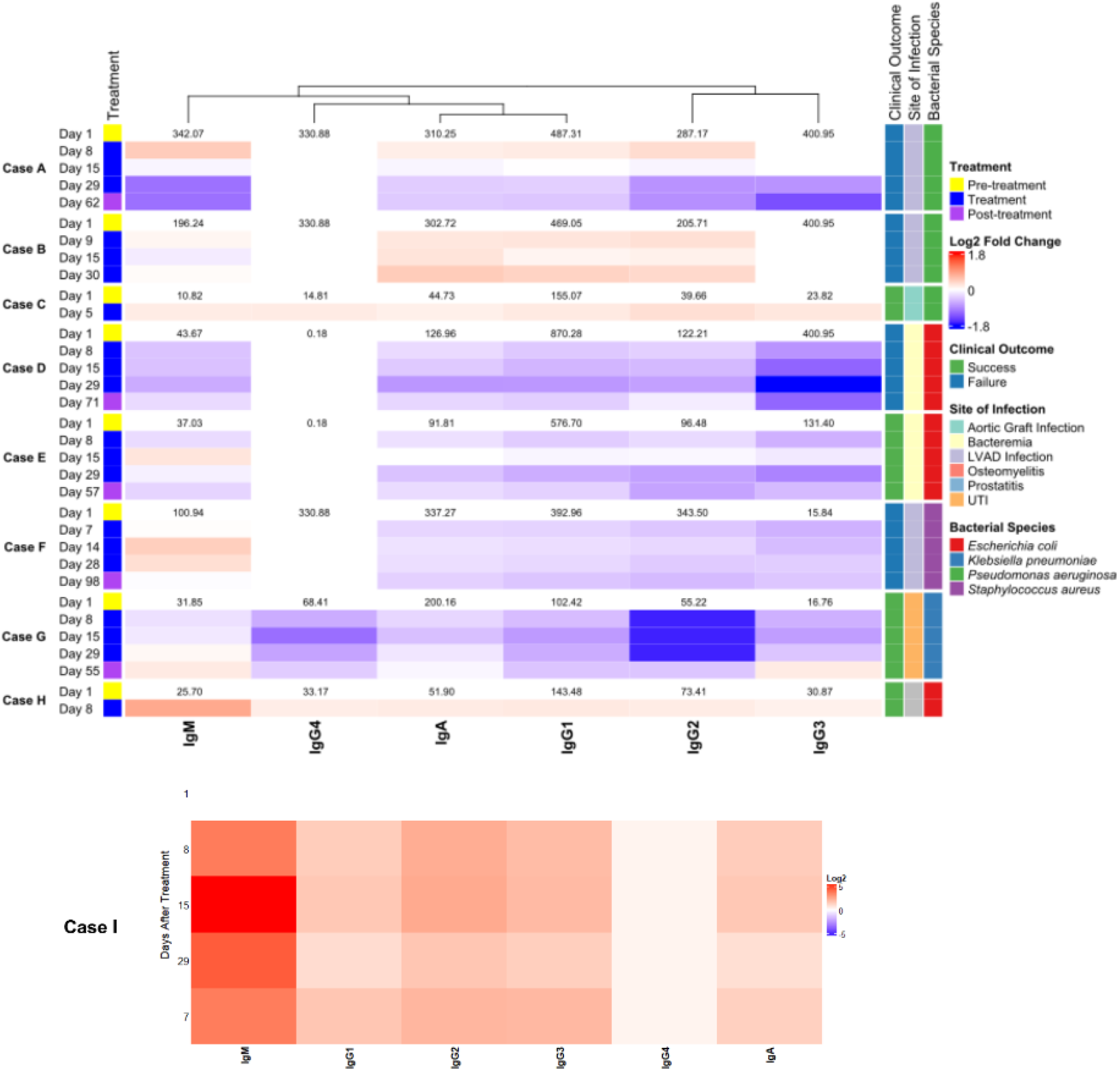
Unified heatmap of serum immunoglobulin levels throughout the course of treatment of all nine cases. This heatmap defines different isotypes and tracks their levels throughout treatment. Additionally, metadata is displayed to the right of the heatmap with a legend. Case I is included separately from the unified heatmap because individual immunoglobulin levels were very high and inclusion with the other cases skewed the heatmap visualization of change for all other cases.

**Figure 6.**
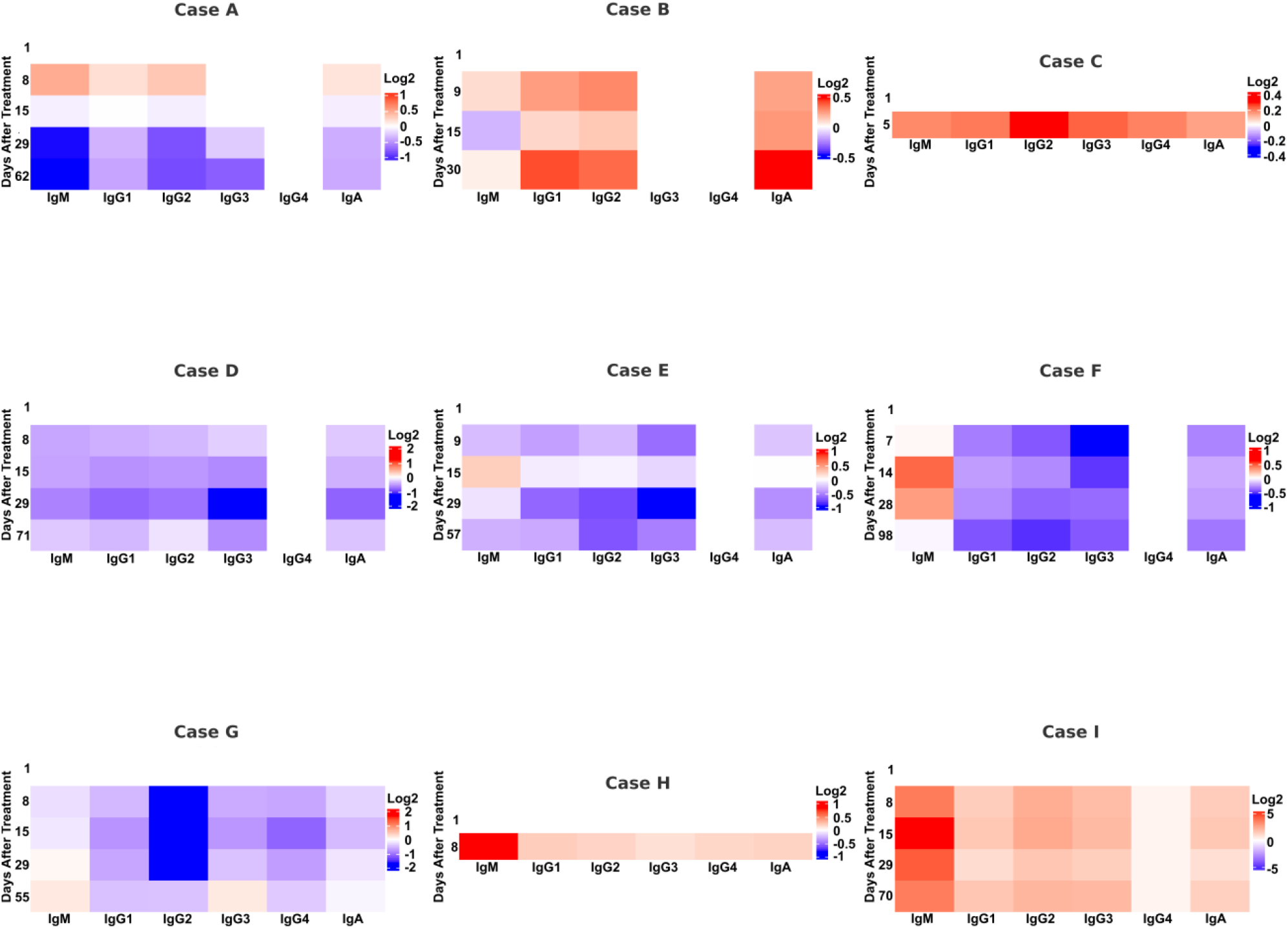
Patient immunoglobulin levels comparing immunocompetent (a) and immunocompromised hosts (b). The IgM response is indicated by a black outlined box. (**a**) Serum samples from immunocompetent cases A, B, F, H, and I were analyzed using an isotyping assay. (**b**) Serum samples from immunocompromised cases D, E, and G were analyzed using an isotyping assay.

Finally, immune responses differed by phage composition. In one patient who received two separate LVAD-targeted *P. aeruginosa* phage regimens, serum neutralization developed against one phage during the first course, necessitating a switch in therapy, whereas the second regimen elicited minimal to no neutralization (Figure 7). These findings suggest variability in phage immunogenicity across preparations or types of phages.

**Figure 7.**
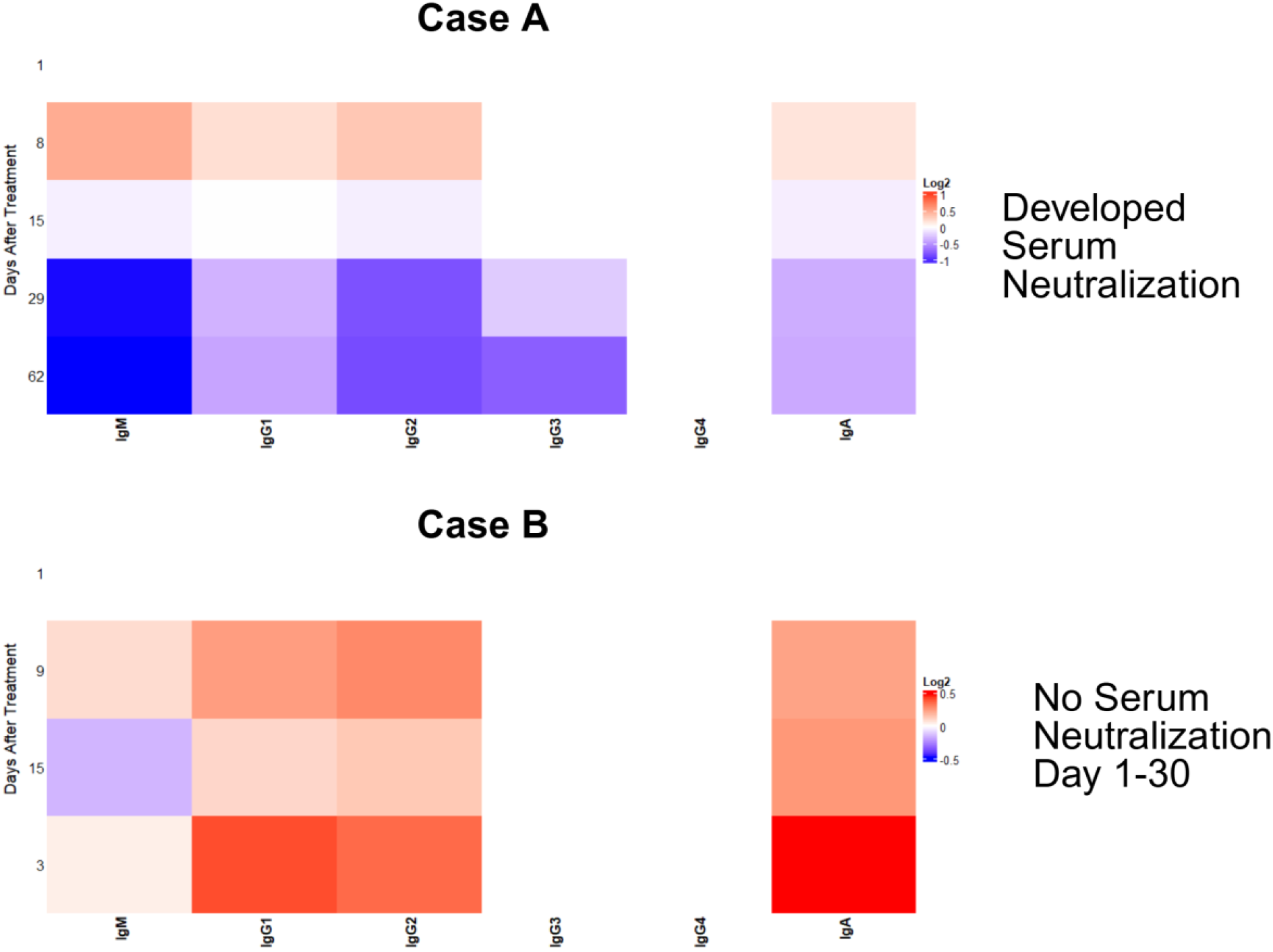
Isotype levels from an LVAD recipient with recurrent *Pseudomonas aeruginosa* bacteremia treated with two separate course of different phages about three months apart. Case A indicates the first round of treatment with this patient. Case B indicates the second round of treatment with this patient, which occurred 3 months after the end of phage therapy in Case A. The first round of treatment developed serum neutralization, and the second treatment developed little to no serum neutralization.

## Discussion

Phage therapy shows promise for the treatment of multidrug-resistant infections, but its interactions with the human immune system remain are not clear. To address this knowledge gap, we analyzed immune responses before, during, and after IV phage therapy in nine compassionate-use cases. Our findings highlight three themes: (1) IV phage therapy commonly elicits an IgM response, particularly in immunocompetent hosts; (2) cytokine responses are variable, influenced more by infection type and host immune status rather than by phage administration itself; and (3) the immune response differs depending on the phages used, suggesting variability in phage immunogenicity.

The transplant recipients that received phage therapy in this series were immunocompromised due to a combination of agents that inhibit both B and T cell function, including agents such as tacrolimus, prednisone and mycophenolate mofetil. Comparisons of the cytokines IL-6, TNF-α, IL-10, and IL-1β between immunocompetent and immunocompromised individuals revealed that immunocompetent individuals had higher levels of these cytokines at all timepoints, including prior to and following phage therapy. However, despite apparent differences between cytokine levels, the clinical outcome (success or failure) of phage therapy was independent of the individual’s immune status. We report that from the five successful cases, two were from immunocompromised hosts, and three were from immunocompetent hosts. Currently, it is thought that synergy with the host immune system may be important for successful phage therapy.^3,8,9^ Supporting evidence is provided from a study on a 7-year-old child with an extensive chronic *P. aeruginosa* osteoarticular infection who received phage therapy.^10^ The phage stimulated the child’s innate and adaptive immune response and contributed to treatment success. However, these claims imply that an immunocompromised individual’s immune system would be unable to cooperate effectively with phage therapy and, therefore, result in unsuccessful treatment.^11^ We introduce contrary evidence; immunocompromised individuals can succeed with phage therapy, and higher levels of various immune system components do not correlate with the outcome of phage therapy.

In 7/9 cases, patients developed an IgM-mediated response within 1-2 weeks of IV phage initiation, consistent with early adaptive immune response. This response was more robust in immunocompetent patients and attenuated in immunocompromised hosts, many of whom were transplant recipients on regimens suppressing both B- and T-cell function. Despite these differences, IgM responses were not associated with treatment success or failure, indicating that antibody development is not associated with clinical outcome.

Cytokine responses were heterogeneous across patients, with no consistent pattern linked to outcome. IL-6, TNF-α, IL-10, and IL-1β levels were higher in immunocompetent compared with immunocompromised hosts, both before and during treatment, but outcome was independent of immune status—two immunocompromised and three immunocompetent patients experienced successful treatment. This observation challenges the view that synergy with host immunity is essential for therapeutic success and adds to prior case reports suggesting that immunocompromised patients may benefit from phage therapy.

When cytokines were analyzed by pathogen, clearer trends emerged. Patients with *P. aeruginosa* infection exhibited marked increases in pro-inflammatory cytokines, including MCP-3 and TNF-β, consistent with prior literature.^12,13^ In contrast, the single *S. aureus* case showed an overall decrease in many pro-inflammatory cytokines, including IL-8, aligning with reports that immune dysregulation in *S. aureus* infections is associated with poor outcomes.^14,15^ These findings suggest that bacterial species, rather than phage exposure itself, may be a stronger determinant of cytokine dynamics during therapy.

We also observed that phage composition influenced the immune response. One patient received two distinct phage cocktails for the same LVAD infection, separated by three months. The first regimen induced a strong IgM response and serum neutralization of one phage by day 11, while the second regimen produced minimal IgM response and minimal to no serum neutralization, despite being directed against the same pathogen. Animal studies support this variability, showing that phage immunogenicity differs by morphotype and even by individual proteins on the phage capsid.^16,17^ This case underscores that phage-specific factors likely shape immunogenicity in humans, with potential implications for phage selection and engineering when developing phage therapy regimens.

Taken together, these findings indicate that cytokine responses during phage therapy are more reflective of host immune status and pathogen type than of phage administration, while humoral responses—primarily IgM—occur commonly but do not determine outcome. Notably, serum neutralization of phages, did not correlate with clinical failure, consistent with reports that phage activity may persist despite antibody formation, possibly through compartmentalization or local delivery or perhaps a shorter duration of phage therapy may have been sufficient.

This exploratory study has limitations, including small sample size, heterogeneity of infections and pathogens, and variable treatment regimens. However, strengths include serial sampling from all patients, consistent IV phage administration in all patients, and paired baseline and longitudinal immune data. To our knowledge, this is the first systematic effort to examine cytokine and immunoglobulin responses in human IV phage therapy.

In summary, we report that (1) most patients mount an IgM response to IV phage therapy within one to two weeks, (2) cytokine responses during phage therapy vary by host immune status and infecting organism and do not impact clinical outcome, and (3) phage-specific factors influence immunogenicity. These results suggest that detailed immune profiling in future clinical trials may help identify patterns of beneficial or detrimental responses. Screening phages for immunogenicity, both in terms of synergy with host immunity and risk of neutralization, may be valuable in optimizing therapeutic design. Further work is needed in larger, more homogeneous populations to define how the human immune response influences phage therapy efficacy and to establish best practices for clinical implementation.

## Author Contributions

Conceptualization, S.R.R, S.A., A.M.; methodology, S.R.R., J.C., K.S., S.A., A.M.; formal analysis, S.R.R., J.C., K.S., S.A.; investigation, S.R.R., J.C., K.S., S.A.; data curation, S.R.R., J.C.; writing—original, draft preparation, S.R.R, S.A.; writing—review and editing, S.R.R., J.C., K.S., S.A., A.M.; project administration, S.A., A.M.; and funding acquisition S.A., A.M.

## Data Sharing

De-identified participant data will be made available upon request to the corresponding authors (S.A., or A.M.).

## Acknowledgements

We would like to thank the Texas Medical Center Digestive Diseases Center and Sridevi Devaraj and Xinpu Chen for their help with the Luminex assays.

## SUPPLEMENT

**Table S1.** Clinical details regarding seven patients treated with nine courses of compassionate use phage therapy. LVAD is left ventricular assist device, IV is intravenous, ESBL is extended spectrum beta lactamase, UTI is urinary tract infection. All Day1 serum samples were obtained at baseline prior to phage initiation.

| Case Name | Immune Status | Diagnosis | Species | Sample Collection Day | Phage Therapy | Clinical Outcome |
| --- | --- | --- | --- | --- | --- | --- |
| A* | Immunocompetent | LVAD infection | *Pseudomonas aeruginosa* | 1 | 50 days of IV phage, plus 1 interoperative dose (Phage from Roach Lab, San Diego State University) | Failure |
|  |  |  |  | 8 |  |  |
|  |  |  |  | 15 |  |  |
|  |  |  |  | 29 |  |  |
|  |  |  |  | 62 |  |  |
| B* | Immunocompetent | LVAD infection | *Pseudomonas aeruginosa* | 1 | 36 days of IV phage (Phage from Walter Reed Army Institute of Research) | Failure |
|  |  |  |  | 9 |  |  |
|  |  |  |  | 15 |  |  |
|  |  |  |  | 30 |  |  |
| C | Immunocompetent | Aortic Graft Infection | *Pseudomonas aeruginosa* | 1 | 42 days of IV phage (Phage from Walter Reed Army Institute of Research)) | Success |
|  |  |  |  | 5 |  |  |
| D** | Immunocompromised (Kidney transplant) | Recurrent ESBL *E. coli* bacteremia | *Escherichia coli* | 1 | 42 days of IV phage (TAILOR Labs) | Failure |
|  |  |  |  | 8 |  |  |
|  |  |  |  | 15 |  |  |
|  |  |  |  | 29 |  |  |
|  |  |  |  | 71 |  |  |
| E** | Immunocompromised (kidney transplant) | Bacteremia | *Escherichia coli* | 1 | 42 days of IV phage (TAILOR Labs) | Success |
|  |  |  |  | 8 |  |  |
|  |  |  |  | 15 |  |  |
|  |  |  |  | 29 |  |  |
|  |  |  |  | 57 |  |  |
| F | Immunocompetent | LVAD infection | *Staphylococcus aureus* | 1 | 42 days of IV phage (Phage from TAILOR Labs) | Failure |
|  |  |  |  | 7 |  |  |
|  |  |  |  | 14 |  |  |
|  |  |  |  | 28 |  |  |
|  |  |  |  | 98 |  |  |
| G | Immunocompromised (kidney transplant) | Recurrent UTI, and Sepsis | *Klebsiella pneumoniae* | 1 | 42 days of IV phage (Phage from Center for Phage Technology Texas A&M University and Li Lab Monash University) | Success |
|  |  |  |  | 8 |  |  |
|  |  |  |  | 15 |  |  |
|  |  |  |  | 29 |  |  |
|  |  |  |  | 55 |  |  |
| H | Immunocompetent | Prostatitis | *Escherichia coli* | 1 | 14 days of IV phage (Phage from TAILOR Labs) | Success |
|  |  |  |  | 8 |  |  |
| I | Immunocompetent | Pelvic osteomyelitis | *Pseudomonas aeruginosa* | 1 | 42 days of IV phage, plus 1 intraosseous injection (Phage from Adaptive Phage Therapeutics) | Success |
|  |  |  |  | 8 |  |  |
|  |  |  |  | 15 |  |  |
|  |  |  |  | 29 |  |  |
|  |  |  |  | 70 |  |  |

\* Case A and B reflect the treatment of the same patient on two separate occasions. There were
104 days between the treatment of case A and B
\*\* Case D and E reflect the treatment of the same patient on separate occasions. There were
313 days between the treatment of case D and case E.

